# Total infectome investigation of diphtheritic stomatitis in yellow-eyed penguins (*Megadyptes antipodes*) reveals a novel and abundant megrivirus

**DOI:** 10.1101/2023.07.09.548243

**Authors:** Janelle R. Wierenga, Kerri J. Morgan, Harry S. Taylor, Stuart Hunter, Lisa S. Argilla, Trudi Webster, Lauren Lim, Rebecca M. Grimwood, Hendrik Schultz, Fátima Jorge, Mihnea Bostina, Laura Burga, Puawai Swindells-Wallace, Edward C. Holmes, Kate McInnes, Jemma L. Geoghegan

## Abstract

First identified in 2002, diphtheritic stomatitis (DS) is a devastating disease affecting yellow-eyed penguins (*Megadyptes antipodes*, or hoiho in te reo Māori). The disease is associated with oral lesions in chicks and has caused significant morbidity and mortality. DS is widespread among yellow-eyed penguin chicks on mainland New Zealand yet appears to be absent from the subantarctic population. Corynebacterium spp. have previously been suspected as a causative agent yet, due to inconsistent cultures and inconclusive pathogenicity, its role in DS is unclear. Herein, we used a metatranscriptomic approach to identify potential causative agents of DS by revealing the presence and abundance of all viruses, bacteria, fungi and protozoa - together, the infectome. Oral and cloacal swab samples were collected from presymptomatic, symptomatic and recovered chicks along with a control group of healthy adults. Two novel viruses from the Picornaviridae were identified, one of which - yellow-eyed penguin megrivirus - was highly abundant in chicks irrespective of health status but not detected in healthy adults. Tissue from biopsied oral lesions also tested positive for the novel megrivirus upon PCR. We found no overall clustering among bacteria, protozoa and fungi communities at the genus level across samples, although Paraclostridium bifermentans was significantly more abundant in oral microbiota of symptomatic chicks compared to other groups. The detection of a novel and highly abundant megrivirus has sparked a new line of enquiry to investigate its potential association with DS.

## Introduction

Diphtheritic stomatitis (DS) is a disease of yellow-eyed penguin (*Megadyptes antipodes*, or hoiho in te reo Māori) chicks first identified in Aotearoa New Zealand in 2002 (Alley et al., 2004, 2017). Since its initial identification, DS has been typically observed biennially, which is thought to be associated with variation in seasonality or nutritional resources (Alley et al., 2017). Over the past five years, DS has been observed every year with a seemingly increasing incidence. DS causes lesions in the oral cavity that usually affects pre-fledgling chicks, typically less than three to four weeks of age but sometimes up to 10 weeks (Alley et al., 2004, 2017). Without treatment, the lesions can result in significant inflammation and plaque formation, making it difficult to feed. Mortality occurs secondary to starvation or aspiration of plaques.

Over the past two decades, there have been multiple attempts to identify the causative agent of DS. Cultures of diseased lesions have identified corynebacteria, including *Corynebacterium amycolatum* (Alley et al., 2017), as well as a novel species of Corynebacterium (Saunderson et al., 2021) later described as *Corynebacterium megadyptis sp. nov*. (Nouioui et al., 2023). However, the association of Corynebacterium with disease is yet to be confirmed. In addition, Corynebacterium species have been identified in healthy yellow-eyed penguin oral cavities (Alley et al., 2017; Saunderson et al., 2021) and these bacteria have been absent from bacterial cultures from the oral lesions of chicks with DS since 2019 (Wildlife Hospital, Dunedin, unpublished diagnostic results). Due to the suspicion of viral inclusion bodies on histological examination of DS lesions within the oral mucosa, as well as the appearance of lesions and inconsistent bacterial cultures, a viral etiology is hypothesised (Alley et al., 2004, 2017). Indeed, both avipoxviruses and herpesviruses have been inconsistently detected upon PCR but, similarly, their potential role in DS has not been confirmed (Alley et al., 2004, 2017). Various other possible fungal and protozoan pathogens have also been investigated, such as Trichomonas sp., Plasmodium sp. and Aspergillus sp., yet causation has not been substantiated (Alley et al., 2017; Sarker et al., 2021).

Despite the many efforts to investigate DS affecting yellow-eyed penguins over the past 20 years, a comprehensive characterisation of all potential pathogens and their association with DS is yet to be undertaken. In this study, we used a metatranscriptomic (i.e., total RNA sequencing) approach to identify all viruses, bacteria, fungi and protozoa species (i.e. the infectome) associated with DS. In 2021, oral and cloacal swab samples were sourced from yellow-eyed penguins located from across the New Zealand mainland population, including from clinically presymptomatic, symptomatic and recovered chicks, as well as healthy adult controls. This study represents the most extensive molecular investigation into DS to date.

## Methodology

### Ethics

This study was conducted under animal ethics approval MUAEC Protocol 21/42 from Massey University and the New Zealand Department of Conservation Wildlife Act Authority 94843-FAU and 94902-FAU.

### Sample collection

Yellow-eyed penguins from three different regions are monitored annually by the New Zealand Department of Conservation along with various organisations across mainland New Zealand. Samples collected in this study originated from North Otago (GPS -45° 21’ 59.99” S,170° 50’ 59.99” E; n = 13), Otago Peninsula (-45° 51’ 17.99” S, 170° 38’ 59.99” E; n = 16) and The Catlins (-46° 29’ 59.99” S, 169° 29’ 59.99” E; n = 14) during the November to December 2021 breeding season (Figure 1). As per their management protocol, newly hatched chicks were monitored and weighed every two to three days. Chicks showing signs of disease (i.e. dehydration, underweight, lethargy, crusting around the eyes or DS lesions in the oral cavity) were treated at the nest or transferred to the Wildlife Hospital, Dunedin for veterinary treatment. If transferred to hospital, affected chicks were immediately placed in incubators for warming based on age (chicks 1-3 days of age at 32°C and a decrease in temperature as the chicks became older). Standard therapy involved 15mg/kg enrofloxacin PO BID and debridement of the lesions with swabs soaked in 10% chlorhexidine BID for mild DS lesions (DS grade 1 to 2). Broad spectrum antimicrobials of 100mg/kg Amoxiclav (400mg amoxycillin and clavulanic acid 57mg) PO BID was initiated for severe or worsening cases (DS grade 2+), refractory to enrofloxacin and/or if the culture/sensitivity results reported a resistance to enrofloxacin and sensitivity to Amoxyclav. Lactated Ringer’s solution was administered subcutaneously (100ml/kg/day) to chicks with severe DS lesions (DS grade 2+) that were unable to tolerate oral fluids. Meloxicam (5mg/kg PO) was provided for severe DS cases (DS grade 3 to 4) once birds were rehydrated. As chicks generally presented with ileus due to being hypothermic and clinically unwell, food was withheld until chicks clinically stabilized, after which fish slurry was provided via gavage up to five times daily as tolerated.

**Figure 1.**
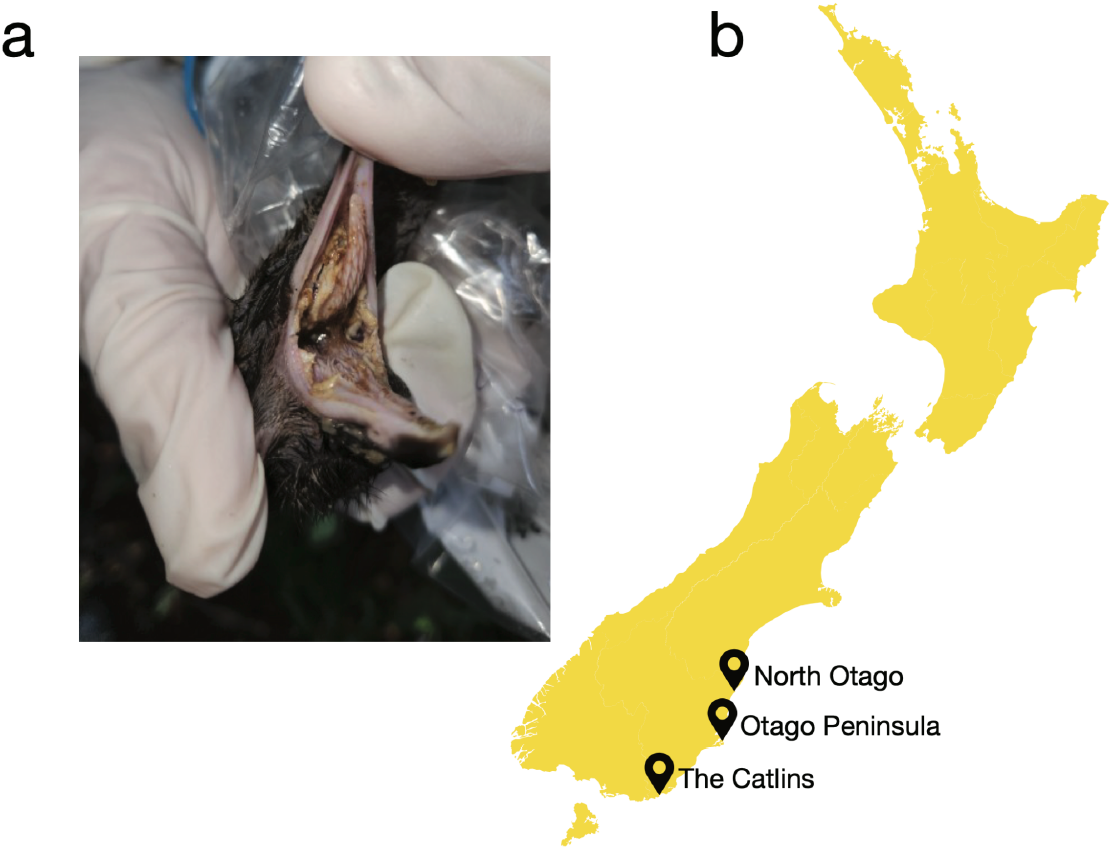
(a) Severe diphtheritic lesions in the oral cavity from a deceased yellow-eyed penguin chick; (b) Sampling locations of the New Zealand mainland population of yellow-eyed penguin chicks suffering from DS.

Oral and cloacal swabs were taken from yellow-eyed penguin chicks at the time of monitoring. Up to three different swab samples were taken from individual chicks until four weeks of age during nest checks. Swabs were categorised based on the health status, location of nest site (North Otago, Otago Peninsula or The Catlins) and type of sample (oral or cloacal swab). Clinical status categorisation included: presymptomatic = chicks appeared healthy with no evidence of DS lesions or dehydration and of appropriate weight, yet developed DS within 1-2 days of sampling; symptomatic = chicks had evidence of DS lesions and were lethargic, dehydrated and underweight; recovered = chicks that previously had DS lesions and were treated with antibiotics with no further evidence of disease; and healthy adults = cloacal swabs taken three months prior with no evidence of disease. Swabs were stored in sterile tubes and immediately placed in DNA/RNA Shield. Tubes were placed into a -20°C freezer within 12 hours of sampling and then transferred to a -80°C freezer within one month of sampling. A random sampling of aerobic and anaerobic cultures along with sensitivities from aerobic organisms from chicks with DS were submitted from the Wildlife Hospital, Dunedin; the decision to culture was up to the veterinarian’s discretion and based on severity of lesions and a random sampling from the three locations. Additional cultures were submitted if chicks were not responding to the standard therapy of debridement and enrofloxacin (noted above).

### Electron microscopy

Oral lesion tissue (n = 2) collected during post-mortem examination in 2022 from chicks with moderate to severe DS lesions were fixed in freshly prepared 2% glutaraldehyde in 0.1M cacodylate buffer pH7.4 for at least 24 hours. The following steps were performed at room temperature under agitation. After primary fixation, tissues were washed three times for 10 minutes with 0.1M cacodylate buffer and post-fixed in 2% osmium tetroxide in 0.1M cacodylate buffer for one hour.

Specimens were then washed three times for 10 minutes with 0.1M cacodylate buffer. Samples were then transferred to a Lynx tissue processor (Australian Biomedical) and run at 20°C under agitation with the following steps: 15 minutes in 0.1M cacodylate buffer, twice in doubled distilled water for 10 minutes, dehydration through a series of ascending grades of ethanol concentrations (50%, 70% and 95%, and 2 x 100% for 10 and 20 minutes), twice with propylene oxide for 15 minutes, followed by two dilutions of Suprr’s resin in propylene oxide (1:1 propylene oxide:resin for 20 minutes and 1:2 propylene oxide:resin for 40 minutes), and finally four Spurr’s resin changes for one hour followed by four-hour incubation time. Each fragment was then transferred to an embedding mould with fresh resin and polymerised for 48 hours at 60°C. Ultrathin sectioning (85 nm) was done with a diamond knife on a Leica EM UC7 ultramicrotome and collected on formvar-coated copper slot grids. Sections were stained with uranyl acetate and lead citrate and investigated using a 120 kV JEOL1400 Flash transmission electron microscope (TEM).

### RNA extraction and sequencing

Swabs were partially thawed and transferred to bead beating sterile vials (VR BashingBead Lysis tubes, 0.1mm and 0.5mm) in DNA/RNA Shield. Samples were lysed in a high-speed bead beater (BioSpec Mini-Beadbeater 24) for a total of five minutes in 1 to 1.5 minute increments with cooling in-between. The supernatant was transferred from the bead beating tubes and proteins, contaminants and enzymes were removed with proteinase K incubation. The samples were then purified, and RNA extracted using the ZymoBIOMICS MagBead RNA kit. RNA was quantified using NanoDrop (ThermoFisher) and RNA from each health status, location and type of sample was pooled into 20 total samples. Extracted RNA was subject to total RNA sequencing. Libraries were prepared using the Illumina Stranded Total RNA Prep with Ribo-Zero Plus (Illumina). Paired-end 150bp sequencing of the RNA libraries was performed on the Illumina NovaSeq 6000 platform.

### Virome composition

Sequencing reads were first quality trimmed then assembled *de novo* using Trinity RNA-Seq (Haas et al., 2013). The assembled contigs were annotated based on similarity searches against the NCBI nucleotide (nt) and non-redundant protein (nr) databases using BLASTn and Diamond (BLASTX) (Buchfink et al., 2015), and an e-value threshold of 1x10^−5^ was used as a cut-off to identify positive matches. We removed non-viral hits including host contigs with similarity to viral sequences (e.g. endogenous viral elements). To reduce the risk of incorrect assignment of viruses to a given library due to index-hoping, those viruses with a read count less than 0.1% of the highest count for that virus among the other libraries were assumed to be contaminants.

### Virus abundance and diversity

Viral abundance was estimated using Trinity with the “align and estimate abundance” tool. The method of abundance estimation used was RNA-Seq by Expectation-Maximisation (RSEM) (Li and Dewey, 2011) and the alignment method used was Bowtie2 with “prep_reference” flag enabled (Langmead and Salzberg, 2012). Estimated viral abundances were first standardised against the number of raw reads in each library and then compared to standardised abundances of ribosomal protein s13 (RPS13), a gene that is seemingly stably expressed in avian hosts (Chapman et al., 2016).

### Virus phylogenetic characterisation

To infer the evolutionary relationships of virus transcripts identified, viral contigs were first translated and then combined with representative protein sequences from the same viral family, which were obtained from NCBI GenBank. All sequences were first aligned using MAFTT (v7.4) (Katoh and Standley, 2013) employing the E-INS-i algorithm and the alignment was trimmed using GBlocks (Castresana, 2000; Talavera and Castresana, 2007) to remove ambiguously aligned regions. Maximum likelihood phylogenetic trees were estimated in IQ-Tree (Lam-Tung et al., 2015) using the best fit amino acid model, LG, as determined by ModelFinder (Kalyaanamoorthy et al., 2017), with 1000 bootstrap replicates. Phylogenetic trees were then annotated in FigTree (http://tree.bio.ed.ac.uk/software/figtree/) and Adobe Illustrator.

### Characterisation of protozoa, fungi and bacteria communities

We characterised total microbial composition differences between health status, swab type and location. Paired-end libraries were first aligned to the *Megadyptes antipodes* genome (GenBank accession: GCA_010078485.1) using the Spliced Transcripts Alignment to a Reference (STAR) software (v2.7.9a) (Dobin et al., 2013) to filter non-host reads. The “outReadsUnmapped” flag was used to keep unmapped (non-host) reads. Filtered reads were then classified using the k-mer-based taxonomic classifier, Kraken2 (v2.1.2) (Wood et al., 2019). A custom Kraken2 nucleotide database was built to include all archaea, protozoa, fungi and bacterial genomes. To assess clustering of microbial communities present in yellow-eyed penguin oral and cloacal swab samples, Kraken2 count data was filtered to genus level. Non-metric multidimensional scaling (NMDS) plots were created using the filtered count data imported into R (RStudio 2022.07.2) (R Core Team, 2021).

Briefly, a distance matrix was created using the vegdist function available in the Vegan package (v2.6-4) (Oksanen et al., 511), with Bray-Curtis dissimilarity as the distance measure. NMDS was performed using the metaMDS function, also available in Vegan, and results were plotted using ggplot (v3.3.6) (Wickham, 2016). The significance of metadata, including swab type, location, and clinical status, on the clustering were tested using a PERMANOVA with the adonis2 function from the Vegan package, with the default usage. Any potential pathogenic bacteria, protozoa and fungi that have previously been associated with disease in avian species were further evaluated (Supplementary Table 1).

### PCR confirmation of a novel megrivirus

Primers were designed to detect and confirm amplicons of yellow-eyed penguin megrivirus using conventional polymerase chain reaction (PCR). Primers were designed using DNASTAR SeqBuilder 12.3.1.48 (DNASTAR) from the complete genome of yellow-eyed penguin megrivirus. In the SeqBuilder Primer Design function, three forward primers and three reverse primers were designed with each primer 25 - 28 bp in length with a GC content of 46.2 - 52.0% and a T^m^ of 58.2 – 63.9°C. Primers were designed and selected based on identifying a product size of 380–520 bp with overlap and no repetition (Supplementary Table 2). RNA was converted to cDNA using a High-Capacity cDNA Reverse Transcriptase kit (Applied Biosystems, ThermoFisher Scientific). PCR amplification was performed with Hot StarTaq Master Mix (Qiagen) as follows: 15 minutes at 95°C; 35 cycles of 94°C for 30 seconds, 57°C for 30 seconds and 72°C for 45 seconds; 72°C for 10 minutes. Optimisation and confirmation included utilising samples with high abundances of yellow-eyed penguin megrivirus obtained from RNA sequencing and by altering the annealing temperature by 1°C to maximise specificity.

Oral lesion tissues were collected during post-mortem examination from six chicks with moderate to severe DS lesions in 2022. Frozen tissue was partially thawed and submerged in lysis buffer containing 1% ß-mercaptoethanol and 0.5% Reagent DX (Qiagen) before tissues were homogenised with TissueRupture (Qiagen). The homogenate was centrifuged to remove any potential tissue residues, and RNA from the clear supernatant was extracted using the Qiagen RNeasy Plus Mini Kit. RNA was quantified using NanoDrop (ThermoFisher).

PCR products were separated by gel electrophoresis with fluorophore (SYBR Safe DNA Gel Stain, Invitrogen, ThermoFisher Scientific) in a 1% agarose gel (Invitrogen, Life Technologies) on 70 mV for 40-50 minutes, and included negative controls, positive controls and 1 kb ladder (Invitrogen, ThermoFisher Scientific) on 0.5X TBE running buffer. PCR bands were visualised by UV light using GelDoc Go Imaging System (Bio-Rad Laboratories). Bands of the expected size were excised, and DNA was purified using a gel cleanup kit (Qiagen QIAquick PCR and Gel Cleanup Kit). DNA was sent for bi-directional Sanger sequencing to the University of Otago Genetic Analysis Services, New Zealand. Forward and reverse sequences were pair-end assembled using Geneious Prime (version 11.0.14, Biomatters Ltd). Contigs were matched to known sequences on the NCBI database using the nt database with Basic Local Alignment Search Tool for nucleotides (BLASTn) (Altschul et al., 1990). Contigs were confirmed with the highest hit as megrivirus. The remainder of the samples of oral lesions were tested individually by converting RNA to cDNA using a High-Capacity cDNA Reverse Transcriptase kit (Applied Biosystems, ThermoFisher Scientific) as noted above, followed by PCR using YEP MGV P1-F and YEP MGV P1-R primers (Supplementary Table 2).

## Results

### Disease characterisation

Between 29 October and 2 December 2021, at least 74% of all recently hatched yellow-eyed penguin chicks (73.7% (n=56/76) of North Otago, 68.6% (n=66/96) of the Otago Peninsula and 82.3% (n=65/79) of The Catlins) demonstrated clinical evidence of DS and were treated as described above. As most nests were monitored frequently, the majority of the chicks were treated as soon as lesions were identified such that DS lesions were mild. Chicks from nests that were monitored less frequently tended to develop more severe DS lesions due to delays in initiating treatment.

Aerobic bacterial cultures from oral lesions were taken by the Wildlife Hospital, Dunedin, New Zealand, from a select group (n = 9) that presented to the hospital (unpublished diagnostic results). The most common organisms cultured were *Clostridium perfringens* and *Enterobacter cloacae*, while other organisms cultured included *Vagococcus lutrae, Staphylococcus sciuri, Clostridium baratii, Paraclostridium bifermantans* (formerly *Clostridium bifermentans*), *Enterococcus faecalis* and *Edwardsiella tarda*. Sensitivities were also performed on the organisms cultured with ∼33% (n = 3) of the organisms resistant to the commonly used antibiotic, enrofloxacin.

### Characterizing the oral and cloacal virome of yellow-eyed penguins

To identify potential pathogens associated with DS, total RNA extracted from swab samples were pooled into 20 libraries based on clinical DS status (presymptomatic, symptomatic, recovered chicks, as well as healthy adults), sample type (oral or cloacal swab) and location (North Otago, Otago Peninsula, The Catlins). Metatranscriptomic sequencing from oral and cloacal swabs yielded libraries containing between 42.2 – 71.6 million paired-end reads that were *de novo* assembled into 677,595 – 1,860,844 contigs.

All 20 libraries were evaluated for viral transcripts. Using a sequence similarity-based approach, exogenous viral transcripts associated with vertebrate hosts were identified using both sequence similarity and phylogenetic analysis (see later), with viral transcripts belonging to the family *Picornaviridae* found in 14 of the 20 libraries. Other viral transcripts identified were thought to be likely dietary (fish) related (families *Picobirnaviridae, Partitiviridae, Astroviridae* and *Caliciviridae*) or associated with invertebrate or bacterial hosts (*Mitoviridae* and *Leviviridae*) since they shared sequence similarity to viruses infecting non-avian hosts.

Novel virus transcripts that shared sequence similarity to viruses within the *Picornaviridae* were identified in both oral and cloacal chick samples from all locations irrespective of health status, while they were absent from healthy adults. In particular, samples from all locations contained several complete viral genomes that shared 59% amino acid sequence similarity to their closest known genetic relative, Penguin megrivirus, identified in faeces of Adélie penguins (Pygoscelis adeliae) (QQG31481.1) and Weddell seals (Leptonychotes weddellii) (QQG31482.1), as well as to a megrivirus (QDY92330.1) from a red-capped plover (Charadrius ruficapillus). The novel virus identified here, provisionally named yellow-eyed penguin megrivirus (YEP megrivirus), fell within the relatively diverse genus, Megrivirus (Figure 2). An additional picornavirus species with only partial polyprotein sequences present was identified only in cloacal samples from presymptomatic and symptomatic chicks from The Catlins. This viral species, for which we recovered only ∼1,200 nucleotides, was termed yellow-eyed penguin picornavirus sp. (YEP picornavirus sp.). YEP picornavirus sp. shared 52% amino acid sequence identity with its closest known genetic relative, goose picornavirus (QUS52516.1) that appears to fall within an undefined genus and a sister clade to kobuviruses, saliviruses and sakobuviruses (Figure 2).

**Figure 2.**
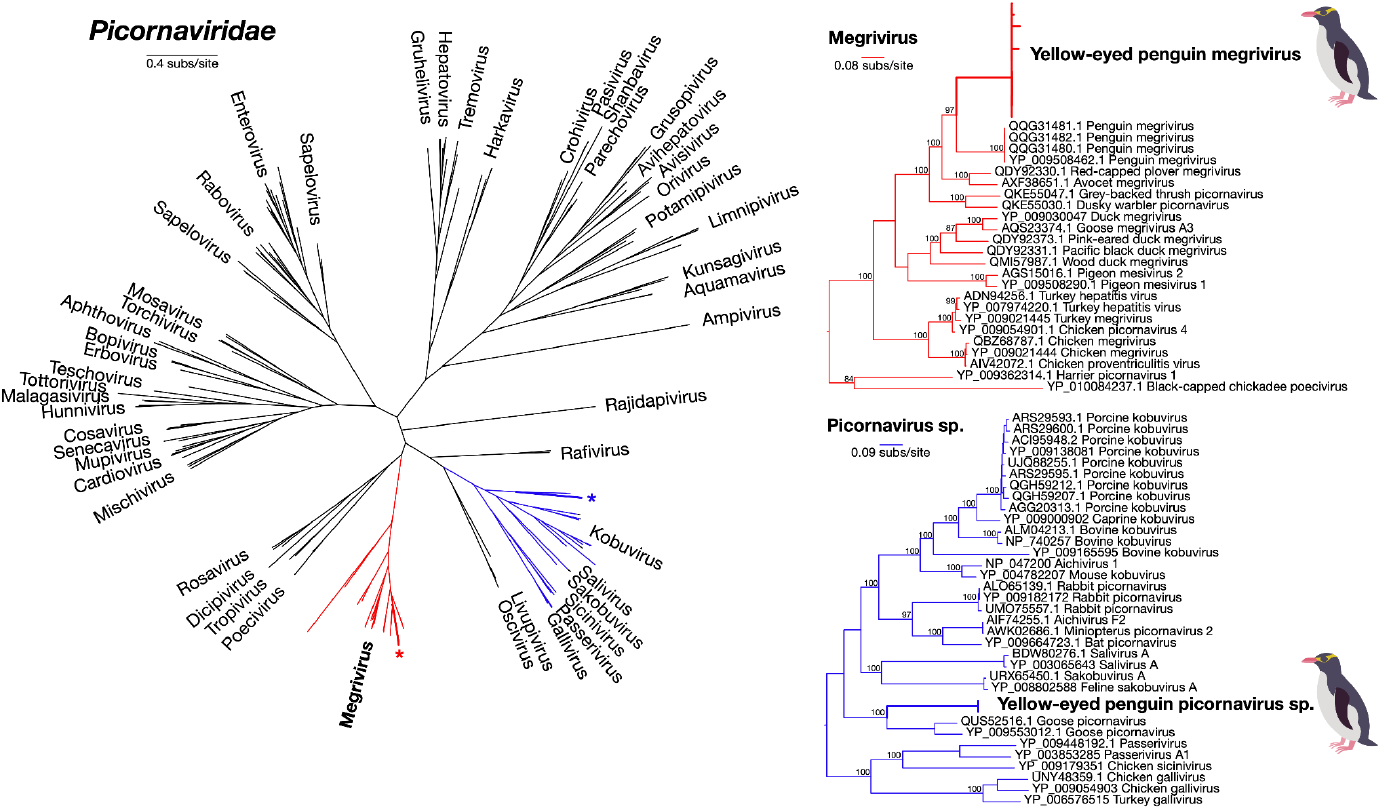
Unrooted maximum likelihood phylogenetic tree (left) showing all currently classified genera from the Picornaviridae. Red branches highlight the Megrivirus genus while blue branches illustrate the closest known relatives of YEP picornavirus sp. The two highlighted clades are shown in more detail (right), illustrating the topological position of the two novel viruses identified (bold). Both clade trees are rooted at their mid-point for clarity and in all phylogenies branch lengths are scaled to the number of amino acid substitutions per site.

Several complete and partial genomes of YEP megrivirus were assembled *de novo*, which all shared >87% amino acid identity within the polyprotein and >90% within the RNA-dependent RNA polymerase. The genome comprised 9,647 nucleotides, containing a single open reading frame for the polyprotein with the structural polyprotein 1 (P1) and the non-structural polyproteins 2 and 3 (P2, P3) (Figure 3). As in other avian megriviruses, the genome organisation contained two putative 2A protein–coding regions and a long three-prime untranslated region (3’ UTR) of 670 nucleotides. The 3’ UTR contained the highly conserved Unit A repeats motif, thought to be common among avian megriviruses (Figure 3) (Boros et al., 2016, 2017).

**Figure 3.**
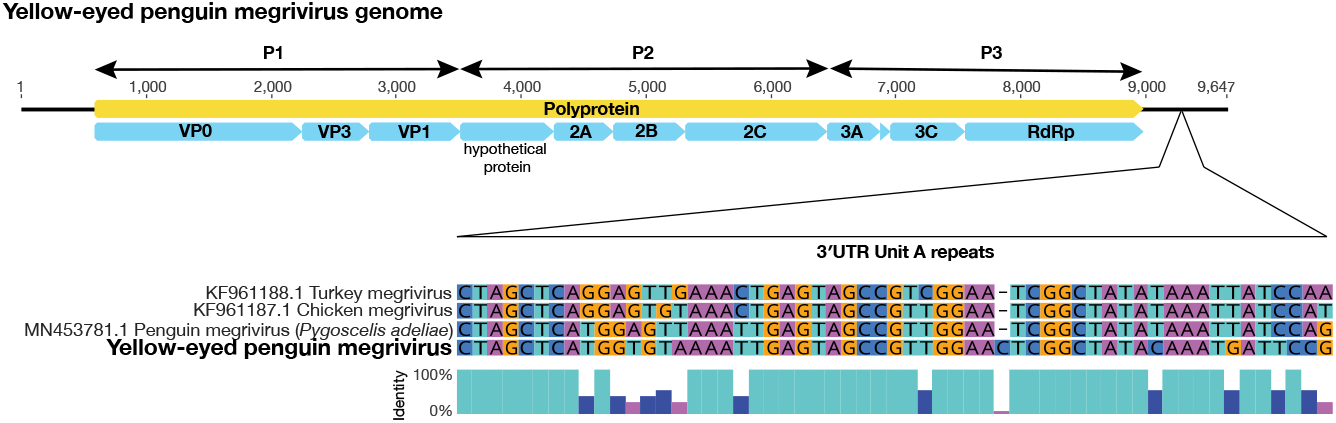
Genome organization of YEP megrivirus (top), showing the positions of the structural polyprotein 1 (P1) and the non-structural polyproteins 2 and 3 (P2, P3). The three-prime untranslated region (3’ UTR), showing the conserved Unit A repeat motif in comparison to penguin, chicken and turkey megriviruses, is indicated within the 670-nucleotide long 3’ UTR section (below).

YEP megrivirus was present in high abundance compared to YEP picornavirus sp. as well as the host gene, Ribosomal Protein S13 (RPS13) (Figure 4). Notably, YEP megrivirus, found in eight of the 20 libraries, was present across all locations and sample types, as well as in presymptomatic, symptomatic and recovered chicks with no overall differences in viral abundance between these groups. In contrast, YEP picornavirus sp., was only found at very low abundances (i.e. several orders of magnitude lower than YEP megrivirus) in cloacal samples from yellow-eyed penguin chicks sampled from The Catlins, in both presymptomatic and symptomatic individuals (Figure 4). No viruses were identified in healthy adults.

**Figure 4.**
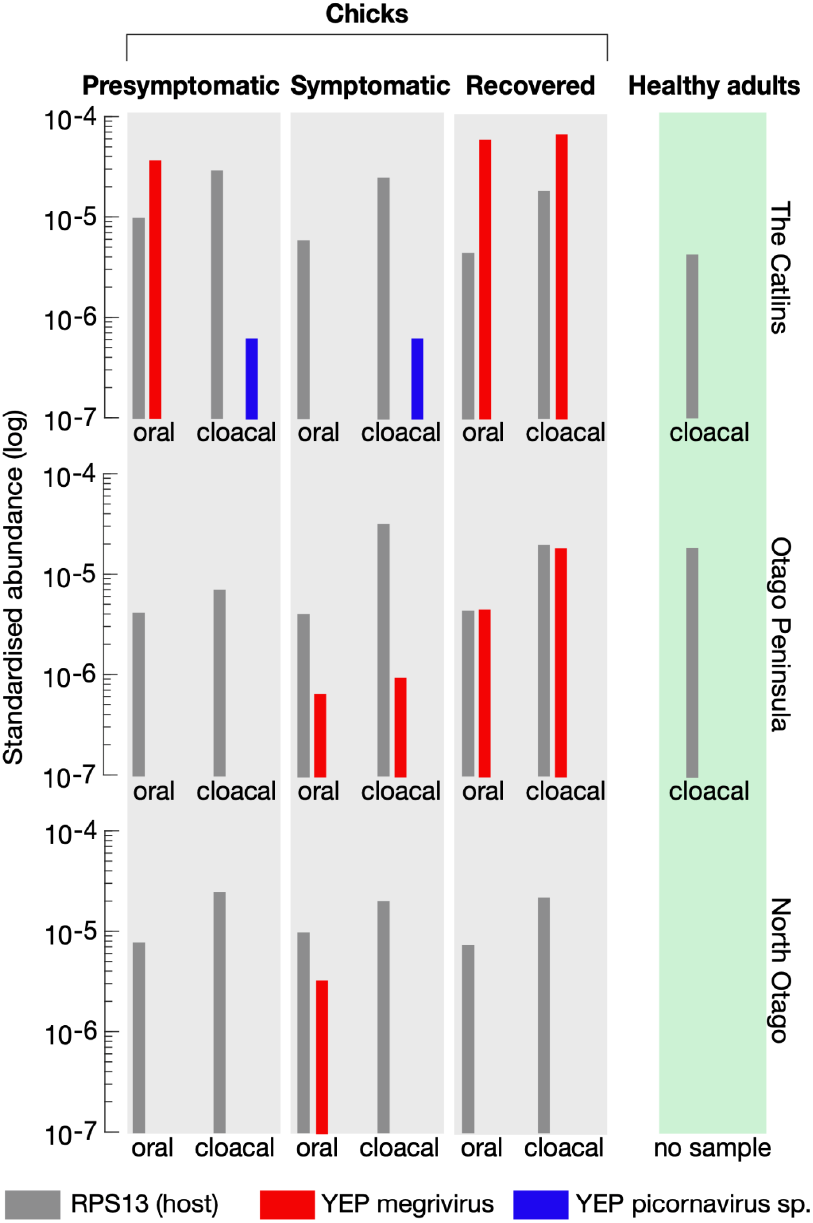
Estimated standardized abundances of two novel virus transcripts within the *Picornaviridae*: megrivirus (red) and picornavirus spp. (blue), from oral and cloacal swabs sampled from three locations. The estimated abundances of megrivirus and picornavirus are shown in each category of presymptomatic, symptomatic and recovered chicks along with cloacal swabs from healthy adults from two locations. Abundances are standardized to the number of paired-end sequencing reads within each library. Estimated abundances of a stably expressed avian host gene, RPS13, is also shown (grey).

### Electron microscopy investigation

Electron microscopy investigation of the oral lesions demonstrated similar cellular remodelling observed in other picornavirus infections (Melia et al., 2019) such as cytoplasmic space filled with multiple vesicles used for viral replication. The samples showed numerous cells exhibiting nuclear condensation, swollen mitochondria and abundant cytoplasmic vesicles (Supplementary Figure 1). Fragmented cells can be occasionally observed inside the tissue. One of the samples was depleted of healthy cells and presented mainly apoptotic cells and high densities of bacteria and nonenveloped filamentous viruses (Supplementary Figure 1).

### PCR confirmation of yellow-eyed penguin megrivirus

PCR was performed to confirm the presence of yellow-eyed penguin megrivirus from tissue samples of DS lesions in the oral cavity. PCR was first optimised using eight swab samples that contained relatively high abundances of yellow-eyed penguin megrivirus sequencing reads. PCR was then performed on individual tissue samples of DS oral lesions from six deceased chicks that were collected in 2022. Four of the six samples tested positive upon PCR for yellow-eyed penguin megrivirus (Supplementary Figure 2). The two negative samples contained low quantities of RNA (<10ng/μL, compared to >100ng/μL for the other four samples), which might explain the negative result.

### Characterising the bacteria, fungal and protozoal communities in yellow-eyed penguins

Non-metric multidimensional scaling ordination was used to characterise both the presence and abundance of microbial communities across all samples, including bacteria, fungi and protozoa, to genus level. As expected, the overall composition of microbial communities differed significantly between oral and cloacal samples of both bacterial (p = 0.001) and eukaryotic genera (p = 0.001) (Figure 5). Notably, neither health status (bacteria: p = 0.63; eukaryotic: p = 0.59) nor sampling location (bacteria: p = 0.96; eukaryotic: p = 0.61) appeared to affect the composition of these microbiotas (Figure 5).

**Figure 5.**
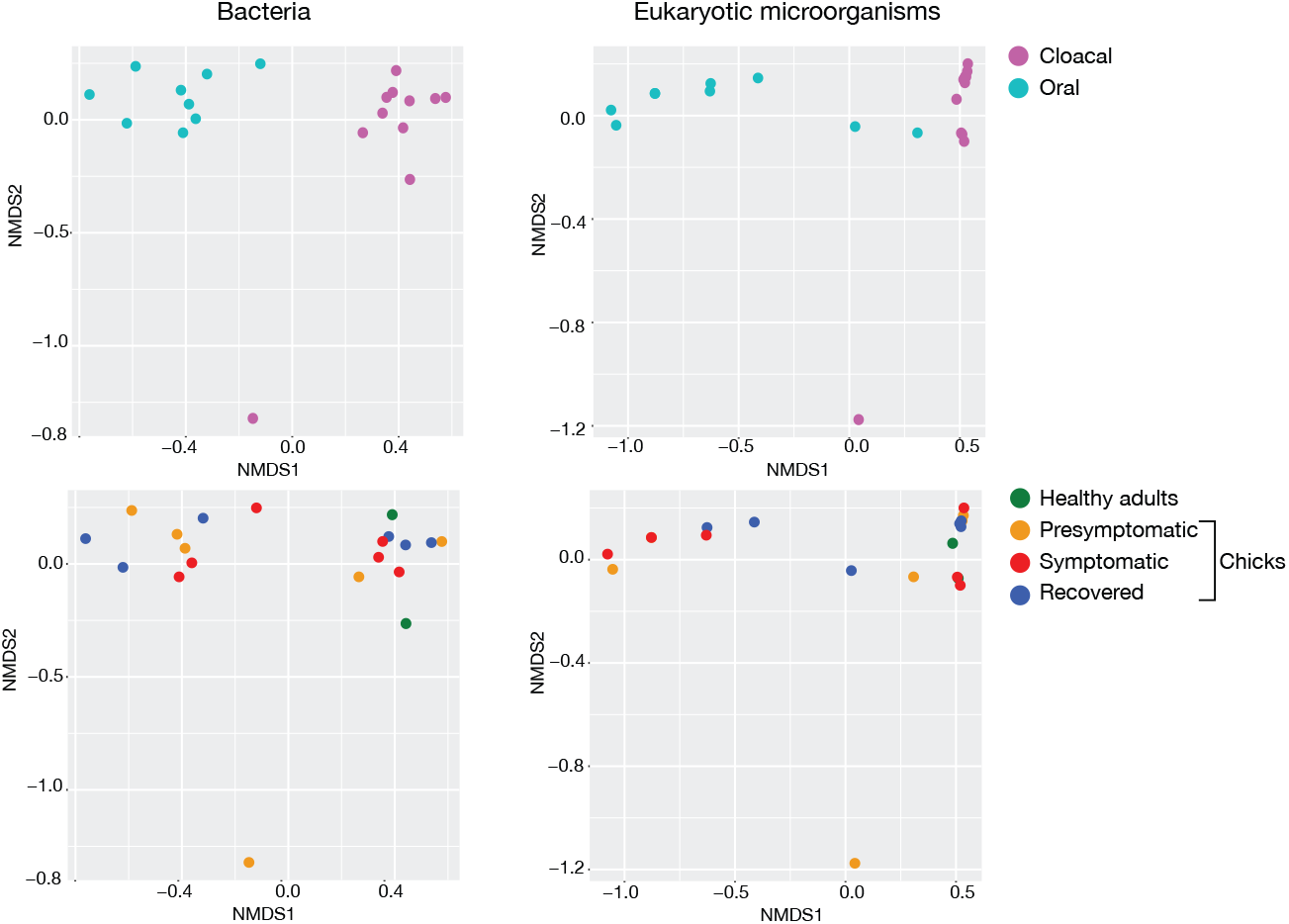
Multidimensional scaling (NMDS) plots showing the effect of sample type (top) and health status (lower) on microbial communities, including both bacterial (left) and eukaryotic microorganisms (right). The full list of microbial communities is available in Supplementary Table s1.

### Potential non-viral pathogens in oral and cloacal samples

Bacteria, fungi and protozoa species that have previously been associated with disease in avian hosts were further analysed (Figure 6). Notably, both *Escherichia coli* in cloacal samples and *Paraclostridium bifermentans* in oral samples from symptomatic chicks were significantly more abundant compared to presymptomatic and recovered chicks, as well as healthy adults (p = 0.05 and p = 0.05, respectively) (Figure 6). However, the difference in *Escherichia coli* abundance was driven by just one sample. *Acinetobacter seifertii* and *Enterobacter cloacae* were slightly more abundant in oral compared to cloacal samples (p ≤ 0.001), regardless of health status. There was also a weak increase in abundance of *Clostridium baratii* in oral swabs taken from symptomatic chicks (p = 0.06) compared to other groups.

**Figure 6.**
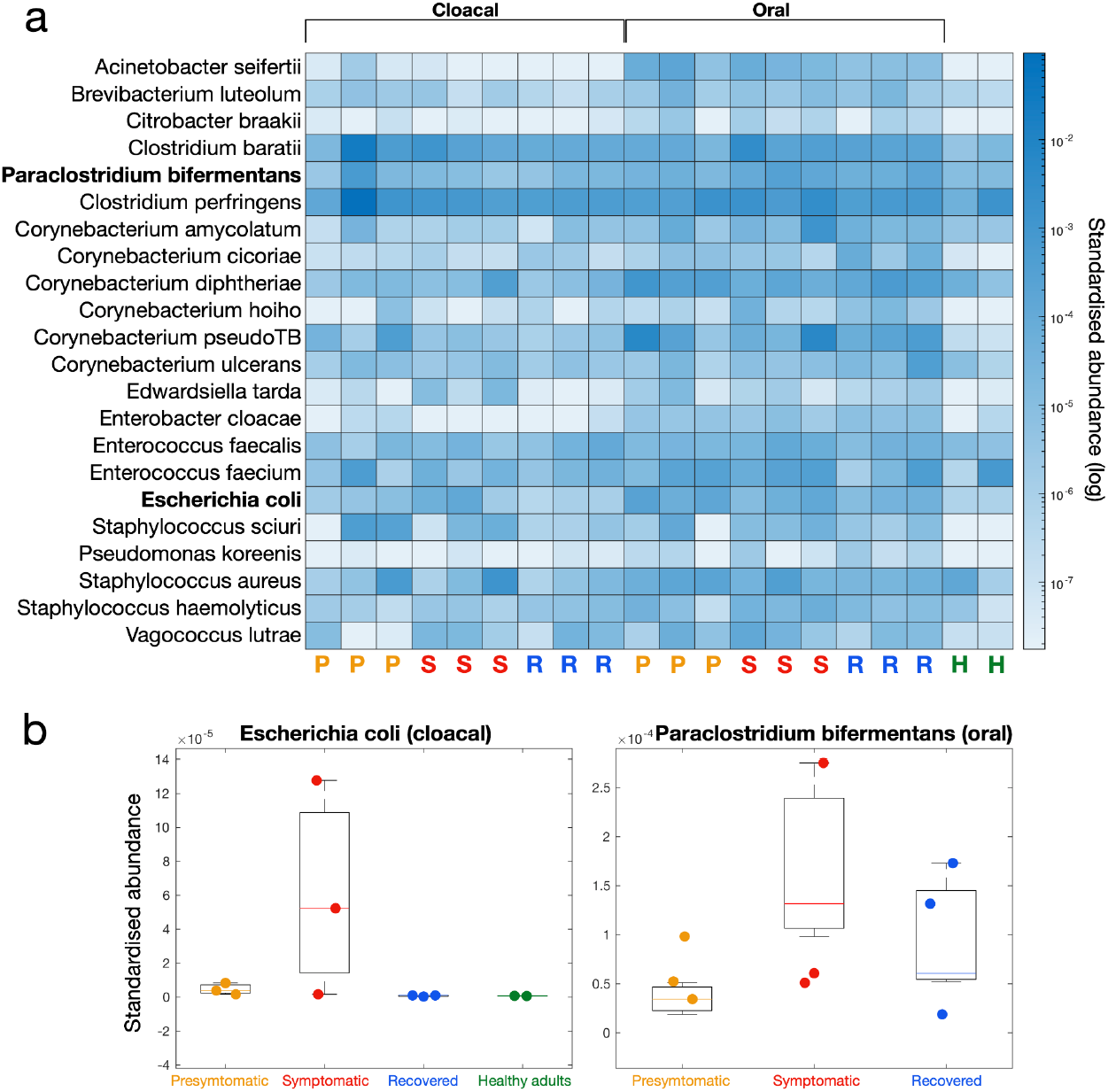
(a) Estimated standardized sequence read abundance of various bacterial species that are known to cause disease in avian hosts. Each column represents cloacal (left) and oral (right) samples from presymptomatic (P), symptomatic (S), recovered (R) or healthy adults (H). (b) Standardized abundance distributions of two bacterial species (*Escherichia coli* and *Paraclostridium bifermentans*) identified where their means significantly differed among symptomatic chicks.

## Discussion

We have described the infectomes of yellow-eyed penguin chicks suffering from diphtheritic stomatitis (DS), including presymptomatic, symptomatic and recovered individuals as well as healthy adult controls. This investigation led to the identification of two novel picornaviruses. One such virus was highly abundant, falling within the genus Megrivirus and termed yellow-eyed penguin megrivirus (YEP megrivirus); the presence of this virus was confirmed by PCR in diseased tissue from biopsied lesions sampled a year later in 2022. Many picornaviruses are known to cause significant disease in both humans and animals, including causing oral lesions (Loeffler and Frosch, 1898; Bachrach, 1968; Ruecker, 1996; Eggers, 1999; Wang et al., 2015). Known avian diseases associated with megriviruses and closely related genera include hepatitis, gastrointestinal disease, encephalomyelitis and diseases of the beak associated with hyperkeratosis (Honkavuori et al., 2011; Boros et al., 2014, 2016; Kim et al., 2015) and, in general, decrease juvenile growth and survival (Honkavuori et al., 2011; Boros et al., 2014). Notably, megriviruses appear to mainly infect birds and have been detected globally in both diseased and seemingly healthy avian hosts (Honkavuori et al., 2011; Phan et al., 2013; Boros et al., 2016, 2017; Zylberberg et al., 2016; Hayer et al., 2021).

YEP megrivirus was detected across all three sample locations. In addition, YEP megrivirus was detected among presymptomatic (one location), symptomatic (two locations) and recovered (two locations) chicks in both oral (n = 5) and cloacal (n = 3) samples, yet it was absent from healthy adult controls. Picornaviruses such as those within the Aphthovirus genus causing foot-and-mouth disease can be persistently shed for prolonged periods upon apparent recovery and can be similarly detected during virus incubation (Brooksby, 1982; Pacheco et al., 2015; Parthiban et al., 2015). In addition, among recovered chicks, the abundance of YEP megrivirus could be artificially inflated due to the recent use of antibiotics depleting bacterial sequencing reads in the sample. Overall, while molecular data alone is unable to determine causation of YEP megrivirus, its detection has sparked a new line of enquiry into its potential association with DS.

We further evaluated the presence of other microorganisms and their role in DS. Bacterial cultures from historical DS lesions over the last two decades most commonly grew Corynebacterium spp. but over the past 10 years, the culturing of bacterial genera was inconsistent, including for Corynebacterium (Alley et al., 2017). Corynebacterium spp. were also cultured from the oral cavities of healthy chicks and adults, little blue penguins and other sea birds (Alley et al., 2017). In addition, the Corynebacterium spp. that were identified were non-pathogenic in an insect challenge model and hypothesised to be opportunistic pathogens (Saunderson et al., 2021). No Corynebacterium spp. were cultured from the 2020-21 and 2021-22 hatching seasons and no evidence of known pathogenic Corynebacterium spp. were found to be associated with disease.

While we found no overall clustering among microbial communities at the genus level across health status, *Paraclostridium bifermentans* was significantly more abundant in oral microbiota of symptomatic chicks compared to other samples. *P. bifermentans* is an anaerobic pathogen that is associated with septic arthritis, osteomyelitis, inflammatory gastrointestinal disease, endocarditis, brain abscesses and lymphadenitis in humans (Kolander, 1989; Edagiz et al., 2015; Hale et al., 2016; Sankar R et al., 2018; Barrett et al., 2020; Zhao et al., 2022). Clostridial diseases have been known to affect avian hosts (Clark et al., 2010; Lueders et al., 2017; Videvall et al., 2020) including penguins ((Rohrer et al., 2023, and unpublished pathology reports), but it is typically only associated with inflammatory gastrointestinal disease and septic peritonitis. Likewise, an increased abundance of *Escherichia coli* in cloacal samples from symptomatic chicks was detected, but only in a single pooled sample. While *E. coli* can be highly pathogenic in birds, they are often also present in the intestinal microflora of healthy birds (Dominick and Jensen, 1984; Leitner and Heller, 1992; Dziva and Stevens, 2008). Further, the majority of diseases associated with avian *E. coli* are secondary to environmental and host predisposing factors (Dho-Moulin and Fairbrother, 1999; Dziva and Stevens, 2008; Collingwood et al., 2014).

In sum, although too preliminary to draw conclusions as to its role in DS, we have identified a novel viral species that warrants further investigation. Assigning pathogen causation in wildlife hosts is challenging, especially in endangered species, but the use of varying paradigms could be useful (Hill, 1965; Evans et al., 1976; Fedak et al., 2015). For example, genomic epidemiological investigations including utilising historical samples collected over the past 20 years might help to identify patterns that support causation. In addition, it is necessary to consider the role of coinfections rather than a single pathogen. Nevertheless, if YEP megrivirus is the cause of DS, it presents a significant concern for the conservation of yellow-eyed penguins in New Zealand.

## Data Availability

The novel viruses can be found under GenBank accession numbers [pending]. Raw sequencing reads can be found [pending].

### Funding

J.R.W. was funded by the Morris Animal Foundation (MAF-D22ZO-418) and J.L.G. was funded by a New Zealand Royal Society Rutherford Discovery Fellowship (RDF-20-UOO-007). J.L.G. was also partially funded by the New Zealand Ministry of Business, Innovation and Employment, Endeavour programme Emerging Aquatic Diseases: a novel diagnostic pipeline and management framework (CAWX2207).

## Acknowledgements

We thank the New Zealand Department of Conservation staff, the Yellow-eyed Penguin Trust, Penguin Place, Penguin Rescue and the Wildlife Hospital, Dunedin. Thanks to Hamish Thompson for the illustration of the yellow-eyed penguin used in this manuscript. Thank you to local Ngāi Tahu members for their collaboration and support.

