## Supplementary Figure 1 for "Total infectome investigation of diphtheritic stomatitis in yellow-eyed penguins (*Megadyptes antipodes*) reveals a novel and abundant megrivirus"

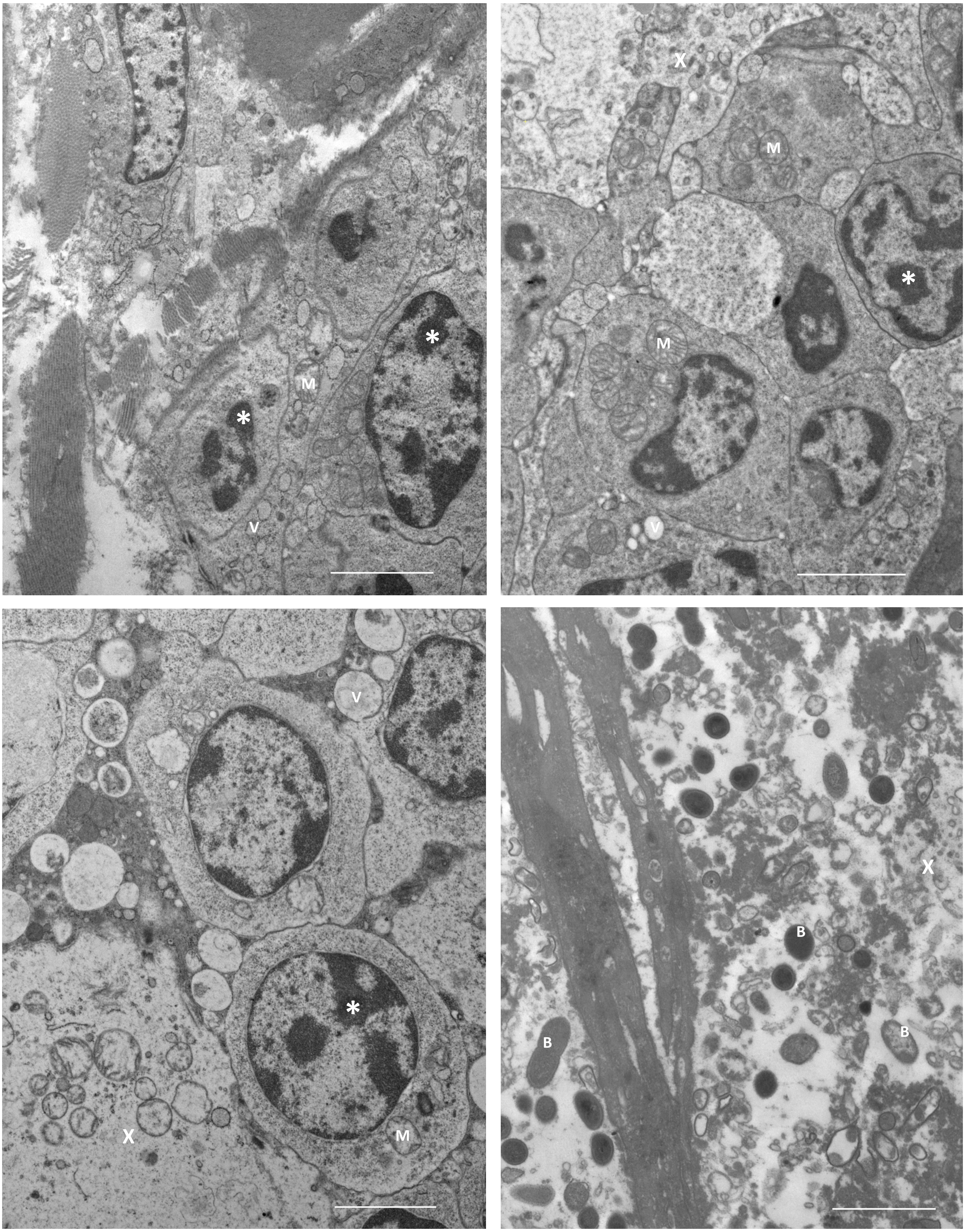


**Supplementary Figure 1.** Transmission electron microscopy of oral lesion tissue. Cells display nuclear condensation (*), swollen mitochondria (M) and abundant cytoplasmic vesicles (V). Fragmented cells (X) and bacteria (B) are occasionally observed. Scale bar 100 nm.
