## Supplementary Figure 2 for "Total infectome investigation of diphtheritic stomatitis in yellow-eyed penguins (*Megadyptes antipodes*) reveals a novel and abundant megrivirus"

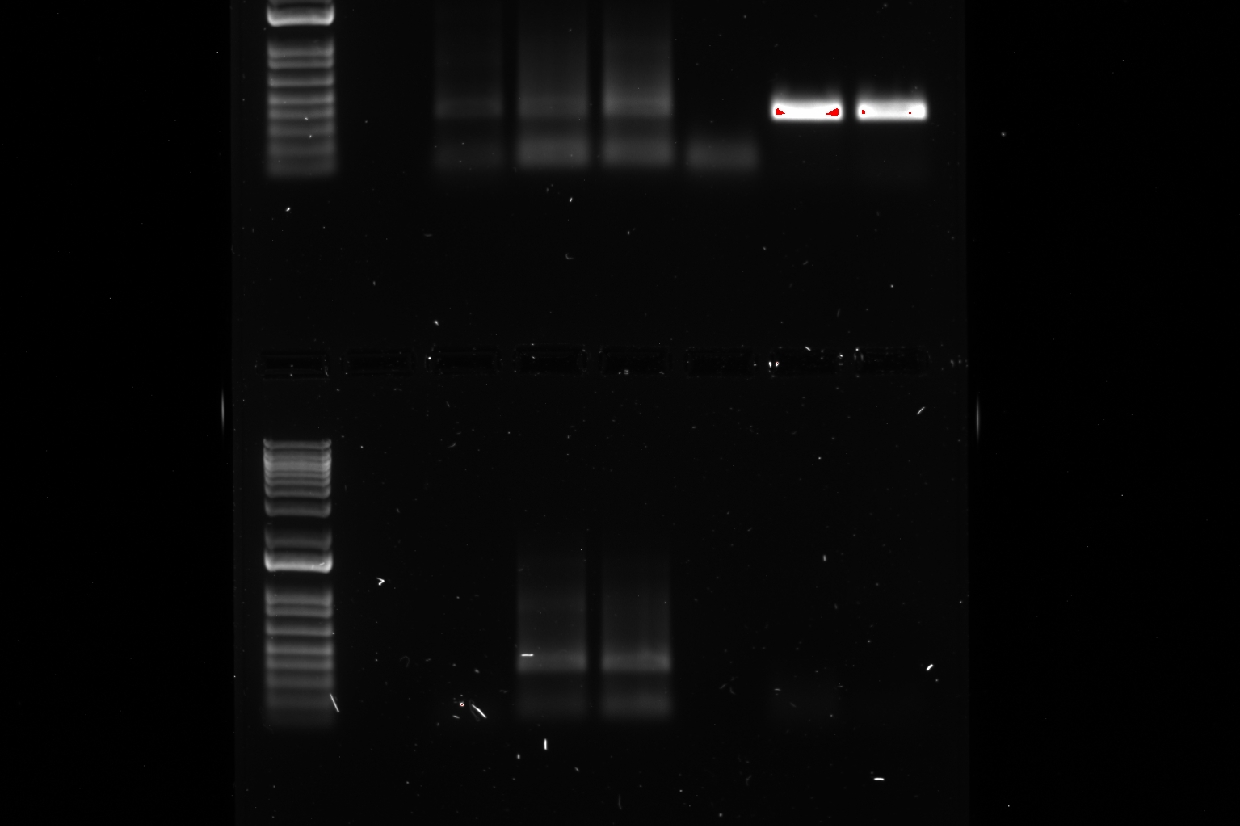

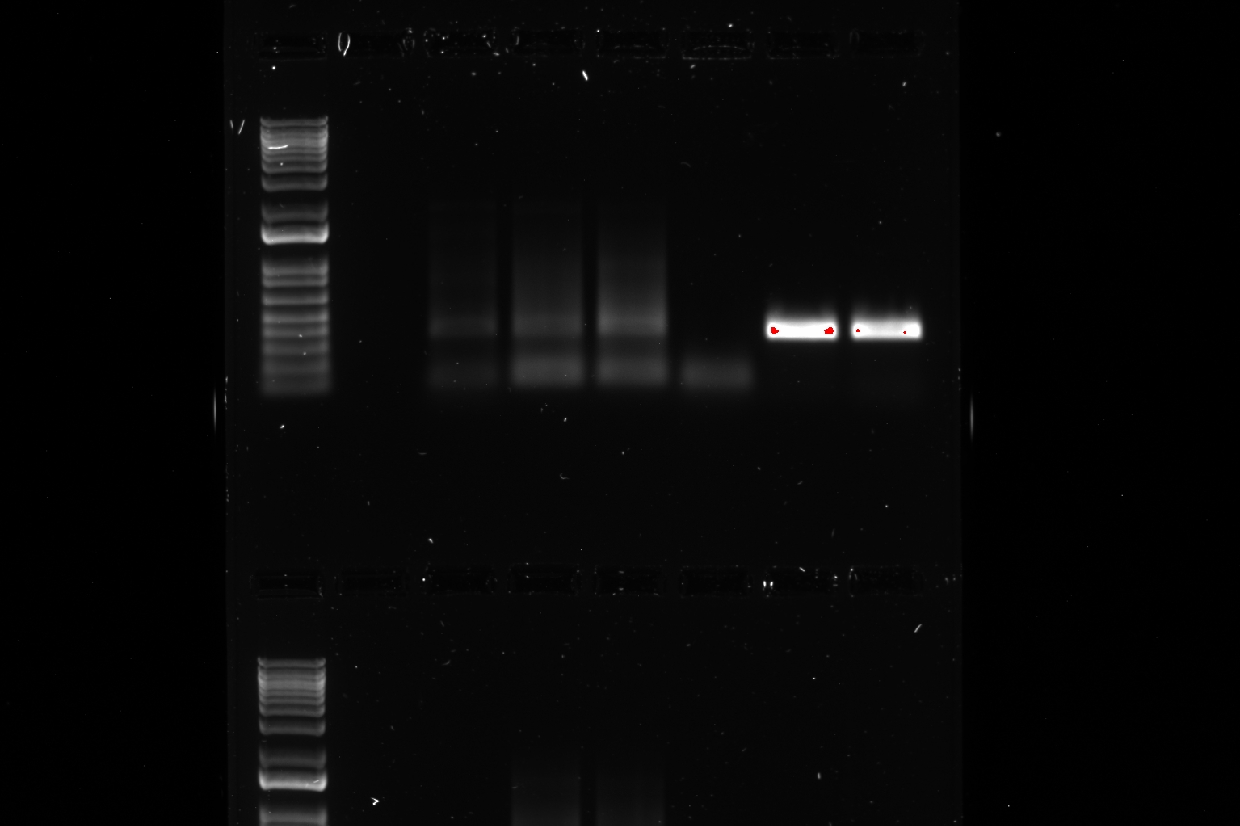

Ld	1	2	3	4	Pos1	Pos2	5	6.1	6.2	Neg

**Supplementary Figure 2**. Gel electrophoresis of extracted RNA from DS lesions with MGV1 primers. Ladder is a 1kb ladder; 1 through 6 are from six chicks with DS lesions evaluated at the time of post-mortem. Chick #6 had severe DS lesions and multiple lesions were collected at the time of post-mortem. Pos1 and Pos2 were positive control samples of oral and cloacal swabs with high abundance of YEP megrivirus. Neg was a negative control sample of DNA/RNA-free water with the same PCR protocol as the positive controls and samples.
