## Supplementary Table 1 for "Total infectome investigation of diphtheritic stomatitis in yellow-eyed penguins (*Megadyptes antipodes*) reveals a novel and abundant megrivirus"

| **Bacteria** | *Acinetobacter seifertii*  *Borrelia spp.* |
| --- | --- |
|  | *Brevibacterium luteolum* |
|  | *Citrobacter braakii* |
|  | *Clostridium baratii* |
|  | *C. perfringens* |
|  | *Corynebacterium amycolatum* |
|  | *C. ciconiae* |
|  | *C. diphtheriae* |
|  | *C. hoiho* |
|  | *C. pseudotuberculosis* |
|  | *C. ulcerans* |
|  | *Edwardsiella tarda* |
|  | *Enterobacter cloacae* |
|  | *Enterococcus faecalis* |
|  | *E. faecium* |
|  | *Escherichia coli* |
|  | *Paraclostridium bifermentans* |
|  | *Pseudomonas koreenis* |
|  | *Staphylococcus aureus* |
|  | *S. haemolyticus* |
|  | *S. sciuri* |
|  | *S. vitulinus* |
|  | *Vagococcus lutrae* |
| **Protozoa** | *Babesia spp.* |
|  | *Cochlosoma spp.* |
|  | *Cryptosporidium spp.* |
|  | *Eimeria spp.* |
|  | *Giardia spp.* |
|  | *Haemoproteus spp.* |
|  | *Hepatozoan spp.* |
|  | *Histomonas spp.* |
|  | *Leucocytozoan spp.* |
|  | *Plasmodium spp.* |
|  | *Toxoplasma spp.* |
|  | *Trichomonas spp.* |
|  | *Trypanosoma spp.* |
| **Fungi** | *Aspergillus spp.* |
|  | *Candidia spp.* |
|  | *Cryptococcus spp.* |
|  | *Dactylaria spp.* |
|  | *Microsporum spp..* |
|  | *Mucor spp.* |
|  | *Rhodotorula spp.* |
|  | *Trichophyton spp.* |

**Supplementary Table 1**. List of pathogenic bacterial, fungal and protozoal organisms evaluated in the 20 libraries from the yellow-eye penguin chick swab samples. The list was compiled from prior disease-causing bacteria, fungi and protozoa in penguins from publications and hospital and MPI culture data (not published) from yellow-eyed penguin outbreak diagnostics.
