## Supplementary Table 2 for "Total infectome investigation of diphtheritic stomatitis in yellow-eyed penguins (*Megadyptes antipodes*) reveals a novel and abundant megrivirus"

| **Primers** | **Nucleotide sequence (5’ 3’)** | **Size (base pairs)** |
| --- | --- | --- |
| YEP MGV P1-F | ACCGATTGGCGAATGTTTGGCAAGG | 380 |
| YEP MGV P1-R | GTTGATCTGCCTGATGGTTGGGTAATCG |  |
| YEP MGV P2-F | TGCATCACCTTTGTCGAGTCAATTGG | 512 |
| YEP MGV P2-R | AAACCCTATACCCCGAACGATAAAGG |  |
| YEP MGV P3-F | CGAATGCCGATGTTATTGGCGTTACAC | 467 |
| YEP MGV P3-R | TGGACGCTTTGATGCTGCAAACACAGC |  |

**Supplementary Table 2**.  Forward and reverse primers designed for YEP megrivirus

PCR testing.
